# Shortcomings of shortcuts: common species cannot fully explain multiscale patterns of species richness

**DOI:** 10.1101/2025.11.11.687838

**Authors:** W. Scott Schwenk

## Abstract

A number of “shortcuts” for assessing biodiversity and prioritizing conservation action have been proposed, one of which is to focus on a subset of the most common species. I critically evaluated the claim that common species better explain patterns of species richness than rare species, using bird data from the same spatial extent at two markedly different spatial resolutions and a third dataset for a larger spatial extent. I did find situations where common species convey more information about species richness patterns than rare species, such as in sequential correlations of species sets with total species richness. However, “hotspots” of species richness tended to be most associated with species occurring with intermediate frequency, rather than the most common (or most rare) species. Furthermore, differences in the degree and sequence of rarity across the three datasets meant that conclusions drawn from species in one dataset did not necessarily hold in the others. Overall, therefore, I found little evidence that common species alone could provide a satisfactory shortcut to understanding biodiversity, particularly given that rare species are often facing the greatest risk of extirpation or extinction. Drawing upon citizen scientists to aid in monitoring and ensuring that unusual or unique ecosystem types and configurations are surveyed may be invaluable in obtaining the thorough understanding of biodiversity needed for successful conservation outcomes.

## Introduction

The sheer complexity and diversity of ecosystems and the species they support has led to the development of a number of “shortcuts” for assessing biodiversity and prioritizing conservation action. One such approach involves identifying one or more surrogate species to represent the needs of other species (Caro and O’Doherty 1999). Many authors have argued that a minority of common or widespread species determine the majority of spatial variation in biodiversity, raising the prospects that such species could serve as simplifying proxies. For example, Lennon et al. (2004) reported that common species are most responsible for richness patterns among British and southern African bird species. Comparable findings have been reported for terrestrial mammals (Vázquez and Gaston 2004), freshwater fish and aquatic insects Cucherousset et al. (2008), butterflies (Pearman and Weber 2007), and vascular plants (Mazaris et al. 2008, Kreft, Sommer, and Barthlott 2006, Pérez-Quesada and Brazeiro 2013). If applicable as a general rule, such a finding has important practical implications. For example, biodiversity monitoring programs could concentrate on a relatively few common species rather than on all species (Mazaris et al. 2008, Landi and Chiarucci 2014). Investigations into the factors responsible for species occurrence could also focus primarily on the most widespread species (Vázquez and Gaston 2004).

Other authors, however, have raised concerns about the reliability of focusing on surrogates for conservation. As emphasized by Rahbek (2005), both the spatial extent and the spatial resolution (grain size) employed can have pronounced effects on resulting patterns of species richness. Hess et al. (2006) found that indicator taxa that perform well at a particular spatial extent and grain may not be effective at different spatial scales and in different regions. The factors that explain species richness also vary depending on spatial extent and grain, with climatic factors commonly most important at large (continental) extents but with environmental diversity, topography and biological interactions among species often increasingly important at more local extents (Rahbek 2005, Whittaker, Willis, and Field 2001). The identification of “hotspots” of species richness – an important practical application of the study of biodiversity patterns – has also been found to be strongly affected by the spatial resolution employed (McKerrow et al. 2018, Shriner, Wilson, and Flather 2006). These findings raise questions about the role of common species in explaining spatial patterns of species richness, their usefulness in identifying species richness hotspots, and their consistency across spatial scales.

In this paper, I critically evaluate the claim that common species better explain patterns of species richness than rare species, using bird data from the same spatial extent at two markedly different spatial resolutions (Vermont Breeding Bird Atlas and Vermont point count dataset) and a third dataset for a larger spatial extent (New York Breeding Bird Atlas). As part of the evaluation, I examine the association of common and rare species with sites of high species richness. I also examine factors associated with individual species occurrence and total species richness among these datasets to better understand the species richness patterns that emerge from looking at accumulated species.

## Methods

### Data sources

#### New York Breeding Bird Atlas

The second New York Breeding Bird Atlas (NY BBA) was a comprehensive survey of the breeding birds of the state of New York conducted from 2000 – 2005 (McGowan and Corwin 2008). The state of New York (141,000 km^2^) was divided into 5,335 5 × 5 km blocks, which were surveyed by more than 1,200 observers who recorded evidence of breeding for species encountered. I eliminated from analysis 295 survey blocks that comprised approximately 50% or less land area within New York due to open water or boundaries that straddled a neighboring state. I expected species occurrence and richness to be substantially reduced in such blocks.

Because limited observer effort per block (number of hours recorded by surveyors) was associated with a reduction in the number of species recorded, I eliminated another 904 blocks with <10 hours of observer effort. The resulting sample (Table 1) consisted of 4,136 blocks (median observer effort = 24 hours). I considered a species to be present if breeding was confirmed, probable, or possible in the block (i.e., “observed” only sightings were eliminated) during any of the survey years.

**Table 1.** Summary of datasets analyzed for this study. Of the original 5,335 study sites (blocks) in the New York Breeding Bird Atlas (BBA), 1,199 were eliminated due to limited observer coverage (<10 h) or limited land area within New York State. Analysis of the Vermont BBA was restricted to sites with the greatest observer coverage (priority 1 and 2 blocks). For the Vermont point count dataset, 24 aquatic or wide-ranging species detected were eliminated from analysis because the study design was not well-suited for assessing their occupancy of study sites.

| Set | Area of sampling unit | # of units | Median observation time per site (hours) | Total # of species observed | Average # (range) of species per site |
| --- | --- | --- | --- | --- | --- |
| NY BBA | 25 km<sup>2</sup> | 4,136 | 24 | 245 | 74.9 (8-129) |
| VT BBA | 25 km<sup>2</sup> | 361 | 60 | 192 | 84.7 (57-121) |
| VT point count | 1.8 ha | 693 | 0.5 | 101 | 10.7 (1-24) |

#### Vermont Breeding Bird Atlas

The second Vermont Breeding Bird Atlas (VT BBA), conducted from 2003 – 2007, closely matched the design of the New York BBA (Renfrew 2013). The state (24,900 km^2^) was divided into 1,237 5 × 5 km blocks that were surveyed by more than 300 observers. I included in the analysis 361 “priority” blocks (Table 1), which were the focus of observer effort and that each received more than 10 hours of observer visitation during the atlas (median = 60 hours). The priority blocks represented a stratified random sample of all blocks (2 blocks out of 6 per USGS 7.5-minute topographic quad). Four priority blocks were eliminated because a significant amount of their land area was outside of Vermont. As in the case of the New York BBA, I considered a species to be present if it was classified as a confirmed, probable, or possible breeder during at least one survey year.

#### Vermont Point Count Dataset

The Vermont point count dataset (Table 1) consists of a stratified random sample of 693 terrestrial points throughout the state, as described in Schwenk and Donovan (2011) and Long et al. (2007). Each point was visited once during the breeding season (20 May to 16 July) in 2003 or 2004, with a visit entailing three separate 10-min counts. Counts were conducted by a single, experienced observer who recorded all bird species heard or seen within 75 m of the point. Because this study was designed to detect and draw inferences about terrestrial species with relatively small home ranges, observations of 24 aquatic or wide-ranging species (such as waterfowl, gulls and raptors) were excluded from analysis.

#### Species richness patterns: common vs. rare species

I examined the role of species rarity in species richness patterns using three approaches. The first approach, following the methods of Lennon et al. (2004), involved sequential correlation of partial species sets. For each dataset, I ranked species from rarest (fewest number of blocks with breeding records or fewest points with detections) to most common. I then generated a series of partial species richness vectors (summed occurrence values at each location) beginning with the rarest species and then sequentially adding the next rarest species until all species were selected. At each step in the sequence, I calculated the correlation between the partial species richness vector and the full species richness vector. I repeated this process in reverse order, beginning with the single most common species and then sequentially adding the next most common species until all species were selected. Finally, I repeated the process using sets in which the order of species was selected randomly. I then compared the correlation results for 10,000 randomly ordered cumulative sequences to the results for the rare-to-common and common-to-rare correlation results for each dataset.

For the second approach, I graphically examined how the degree of rarity of individual species was related to species richness at the sites at which they occurred and at the sites where they were not recorded. I did not perform statistical analyses on these results because the patterns of association between rarity and species richness are mathematically constrained (Arita et al. 2008) in a way that violates standard statistical assumptions.

In the final approach, I tested whether the degree of rarity of species was related to their occurrence in “hotspots” of species richness. I defined “hotspots” as the 10% of locations in each dataset with the most species recorded (Heegaard, Gjerde, and Sætersdal 2013). For each species, I calculated the difference between their frequency of occurrence within hotspots and their frequency of occurrence within the other 90% of locations. I then divided species into quartiles of occurrence frequency and compared mean differences among quartiles within each dataset using analysis of variance.

Environmental and Landscape Factors Associated with Species Occurrence and Richness

#### Individual Species

To further explore patterns of species richness, I investigated factors associated with occurrence of individual species. In the case of the NY and VT BBAs, I used logistic regression to assess five environmental and landscape covariates anticipated to influence species occurrence. Four covariates were the same for both datasets: landscape diversity, road density (included quadratic term), percent wetlands, and elevation. For the New York BBA, a fifth covariate I included was whether the block was located in a coastal / Great Lakes ecoregion (based on seven Atlas-defined ecoregions). For the Vermont BBA, a fifth covariate was latitude. Observer effort (log-transformed hours) was used as a detection covariate in both datasets. Landscape diversity was measured with the Shannon diversity index applied to percentages of land cover classes occurring within blocks. Land cover classes were derived from the 2001 NLCD (MRLC 2001); I consolidated a few similar categories (e.g., grassland and pasture) for analysis purposes. Road density (combined primary, secondary, and local roads) was based on the U.S. Census Bureau 2010 TIGER roads for New York and the E911 roads dataset for Vermont. Percent wetlands was measured as the total percentage of the land area of an Atlas block classified as any type of wetland by the 2001 NLCD. Average elevation of a block was based on the USGS National Elevation Dataset 30 m DEM. The analyses included 180 species for New York and 80 species for Vermont, omitting species with insufficient (<50) occurrences or absences for modeling.

For the VT point count dataset, I summarize the results of single-season occupancy models for 67 species reported by Schwenk and Donovan (2011). The models included six environmental covariates: whether or not the point was primarily forested; an index of topographic wetness (quadratic; larger values represent downslope accumulation of flow); distance to the edge of another major land cover type; percent forest within 1 km (quadratic); percent evergreen forest within 300 m (quadratic); and road density (quadratic) within 1 km of the study point. Differences in the set of covariates I selected for the point count dataset and BBAs reflect the much different spatial scales between the two types.

I assessed relative support for covariates by applying Akaike’s information criterion (AIC) to model sets containing all possible combinations of the covariates (Burnham and Anderson 2002). I report results for covariates occurring in models with a summed AIC weight of greater than 0.5 (out of a possible 1.0), which I define as representing substantial model support.

Because many species were too rare to assess using our modeling approaches, I used qualitative assessment of habitat preferences to better understand factors contributing to rareness and commonness for the complete set of species. Using the Breeding Bird Atlas reports (Renfrew 2013, McGowan and Corwin 2008) and other published sources, I assigned each species to one of six categories of generalized habitat preference: evergreen forests; deciduous and mixed forests; lakes and coastal areas; nonforested uplands; shrublands and young forests; or wetlands. I then reviewed patterns of occurrence of these species within quartiles of species frequency of occurrence.

Finally, I plotted frequency of occurrence for each species in the Vermont point count dataset vs. its frequency in the Vermont BBA, using nonlinear regression to illustrate how rarity related to the spatial resolution considered.

#### Species Richness

To better understand how patterns of species occurrence accumulated into species richness, I examined associations between species richness and sets of environmental factors closely aligned to those used for individual species. For the NY and VT BBAs, I applied multiple regression that accounts for spatial autocorrelation among neighboring blocks (simultaneous autoregressive models with errorsarlm procedure of the spdep package of R 2.15.1 (R Core Team 2014)). As in the case of the individual species logistic regressions, the multiple regressions included landscape diversity, road density (quadratic), percent wetlands, and elevation (quadratic for New York). Each of the seven Atlas-defined ecoregions were also covariates in the NY BBA multiple regression. I predicted that the effect of elevation would differ among ecoregions, so I included interaction terms for the coastal ecoregion and the two high elevation ecoregions (Northern Appalachians and High Alleghenies). Latitude was included as a covariate for the VT BBA. Observer effort was included as a detection covariate for both BBAs.

As with the BBAs, I applied simultaneous autoregressive models to the VT point count Dataset. I used the seven covariates used for the occupancy models for individual species, plus elevation and latitude. I included time of day and date of year as detection covariates as included in the occupancy analysis.

## Results

### Comparison of species occurrence and species richness datasets

Species frequency of occurrence followed a bimodal distribution for both the New York and Vermont BBA datasets (Fig. 1). The New York BBA dataset in particular had a large number of uncommon species: 100 of the 245 species were recorded in fewer than 5% of the blocks analyzed for this study. On average, a species was recorded in 31% of NY BBA blocks (median 12%) compared to 44% of VT BBA blocks (median 38%). For the Vermont BBA, all except one of the 192 species were also recorded in New York. Of the 54 species not recorded in the Vermont BBA, all but three were among the 100 rarest in New York. Approximately half of the species recorded only in New York were restricted primarily to the Atlantic coast, which does not occur in Vermont. Many others were species with ranges that do not extend as far northeast as Vermont. At the opposite end of the rare to common gradient, species recorded with high frequency in one BBA usually were recorded with high frequency in the other. Eight species were recorded in >95% of sites in both datasets.

**Figure 1.**
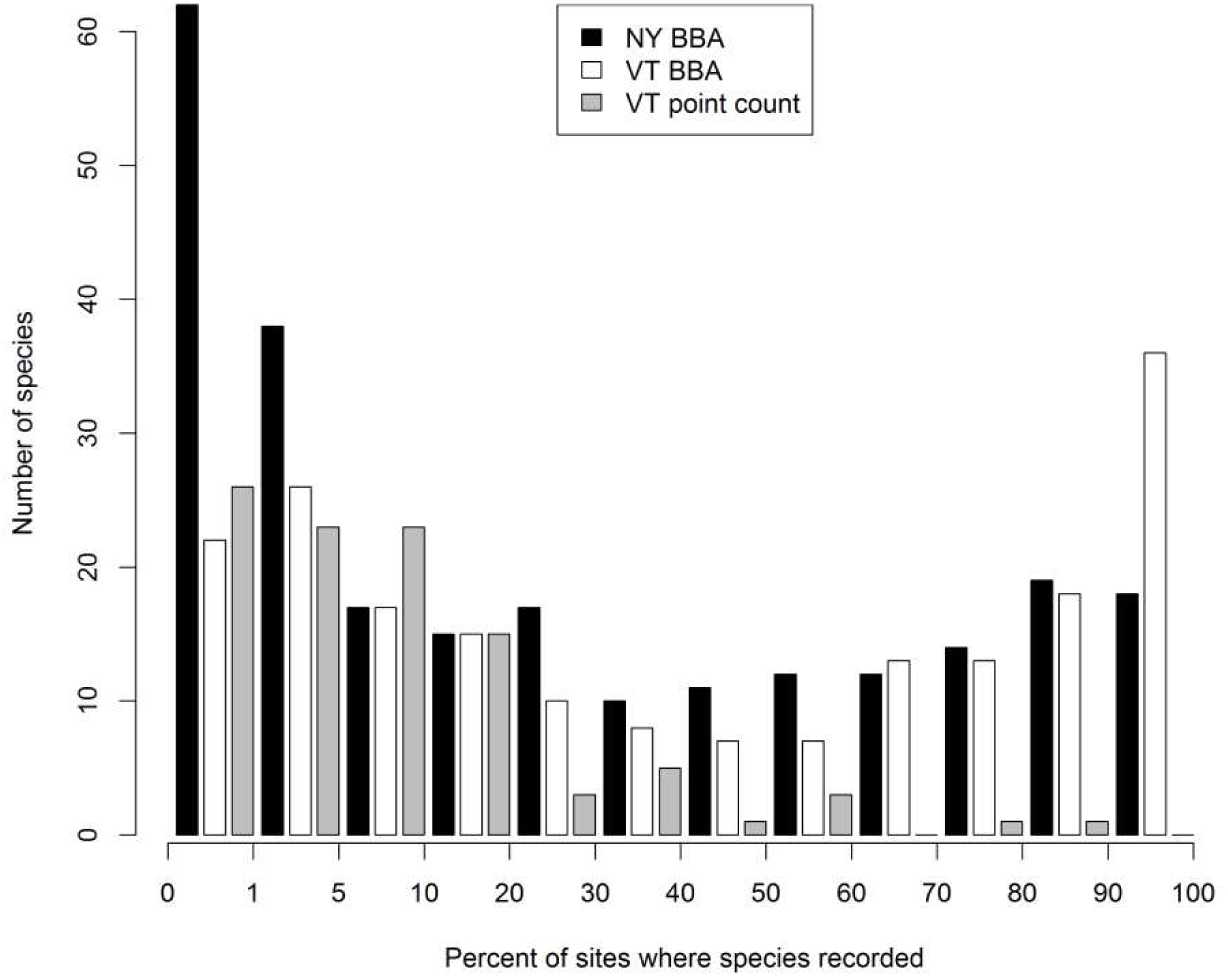
Histogram of frequency of occurrence by species in the New York and Vermont BBAs and the Vermont point count dataset. For display purposes, the horizontal scale of the first three occurrence bins (species recorded in 0-10% of sites) differs from the remaining bins.

The spatial resolution of the datasets influenced the results in a number of ways. The Vermont point count dataset, which used much smaller sampling units over brief time periods, differed from the BBAs in the shape of its distribution for species frequency of occurrence (Fig. 1) and in the inferred degree of rarity of species. Frequency of occurrence was unimodal, with 49 of the 101 species recorded at fewer than 5% of sites. The correlation of frequencies between the Vermont datasets was highly non-linear (Fig. 2); for example, all species occurring with at least 5% frequency in the point count dataset occurred at >50% of BBA blocks. The degree to which a species observed in the point count dataset was correlated with species richness was not highly predictive of whether that species would also be correlated species richness in either the Vermont (*r* = -0.11, based on ranking each species from most to least correlated with richness) or New York (*r* = -0.05) BBA datasets. In contrast, considering the same set of species shared among all three datasets, the Vermont and New York BBA species richness correlations were much more closely related (*r* = 0.73).

**Figure 2.**
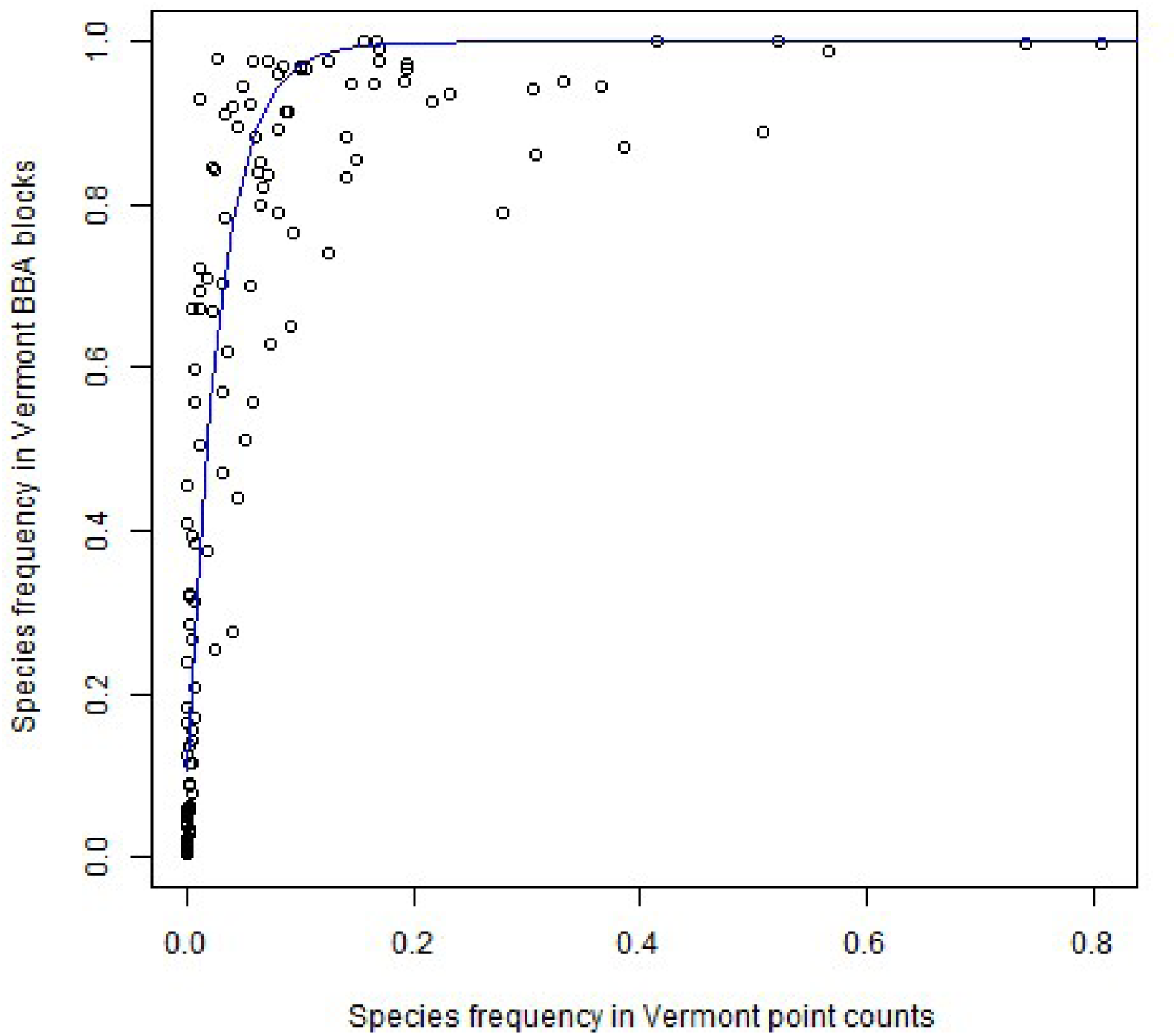
Relationship between recorded frequency in Vermont point counts (1.8 ha, 30 min observation) and Vermont BBA blocks (25 km^2^, >10 hours observation). Each point represents one species. In addition to 101 species shared in the datasets, the figure includes 31 species detected only in the Vermont BBA but judged to have been detectable by the design of the point count surveys. Fitted line is an exponential function (*r* = 0.82).

Average species richness was similar for the BBAs and much lower for the Vermont point count dataset (Table 1). For the blocks analyzed in this study, average species richness was 74.9 for the New York BBA and 84.7 for the Vermont BBA, which also had somewhat higher average observer effort per site. Average species richness was 10.7 for the Vermont point count dataset. Given the small size of the point counts (75 m radius circles), the counts typically encompassed one or at most two major ecosystem types such as grasslands and deciduous forests, providing less opportunity to accumulate species. BBA blocks encompassed a matrix of ecosystem types with differing species.

Although often ignored in analyses of species richness, the degree to which rare species may have been missed in surveys merits consideration. The survey blocks included in this analysis accounted for more than 95% of the species identified as potential breeders anywhere in the respective states of New York and Vermont during the BBAs. The Vermont point count dataset, on the other hand, encompassed a less complete survey of the region’s avifauna. I estimate that based on the Vermont BBA results, approximately 31 additional species of breeding birds (>20% of the total species pool) were available for detection across the state but not observed during the point counts, only counting the types of species for which the point counts are designed. Most of these species were infrequently encountered in the Vermont BBA (<5% of blocks).

### Species richness patterns: common vs. rare species

The first approach to examining whether common species better explained species richness patterns than rare species, which applied sequential correlation of partial species sets, produced qualitatively similar results for the NY BBA, VT BBA, and VT point count datasets (Fig. 3). In all three cases, correlations of cumulative sets with the complete set more rapidly approached unity for common species. This pattern was strongest for the NY BBA, which had the most uncommon species and was the largest dataset in terms of study sites (blocks) and species recorded. The VT BBA, with the most symmetrical distribution of occurrence frequencies, had the least differences between common and rare species sets (Fig. 3).

**Figure 3.**
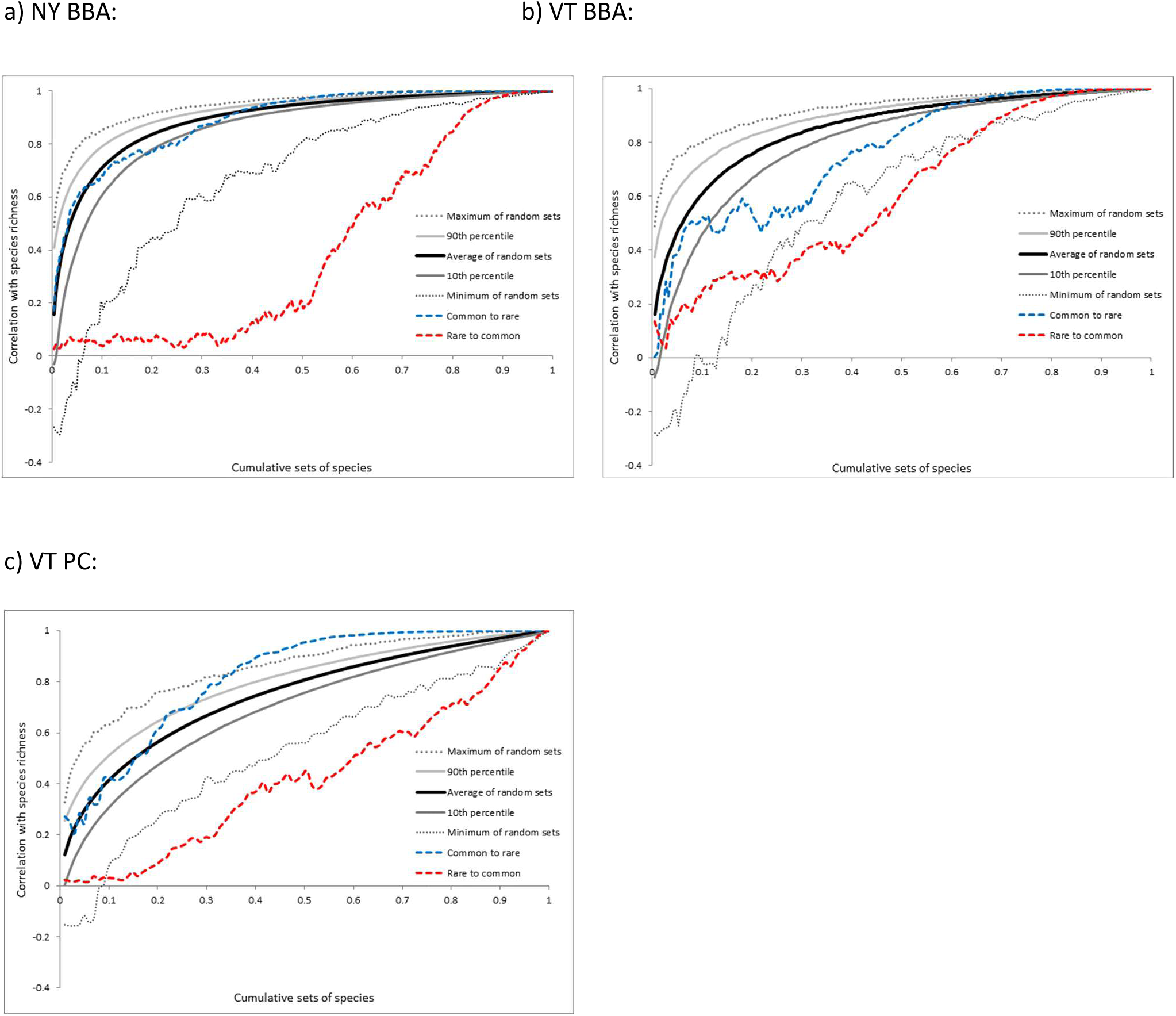
An approach to examining the role of rare and common species in determining species richness patterns: sequential correlation of partial species sets with the full species richness set. Rare to common sets (dashed red lines) were generated by beginning with the site occurrence history of the rarest species and sequentially adding, one species at a time, the next rarest species until all species are included. Common to rare sets (dashed blue lines) are generated the same was except beginning with the most common species and continuing to the rarest species. The remaining lines present summary outcomes for 10,000 iterations of sequential correlations of species selected in random order. Panels: a) New York BBA; b) Vermont BBA; c) Vermont point count dataset.

Consideration of the cumulative sets of randomly selected species (Fig. 3) results in a somewhat more complicated picture than considering only the common-to-rare and rare-to-common sets. Random cumulative subsets almost always converged on unity more quickly than rare-to-common species. Random subsets also frequently approached unity more rapidly than common-to-rare sequences in the case of the Vermont BBA, but for the other two datasets common-to-rare subsets often were more correlated with species richness than were most random subsets. Even for the latter datasets, however, many random sequences were produced that out-performed sequences of the most common species.

Adding results from the second approach provides a fuller, more complex picture of associations between rarity and species richness. Average species richness at sites where rare species were observed was highly variable and provided much more information about species richness than the common species (Fig. 4, left panels). Common species occurred at sites with species richness that, on average, differed little from the dataset means. Arita et al. (2008) and Borregaard and Rahbek (2010) have demonstrated that points in the figure are mathematically constrained to occur only in certain regions; these regions are defined by the dashed lines in Fig. 4. Consider a species that occurs at only one site: it would attain maximum species richness if it occurred at the site with the greatest species richness (which defines the upper line in Fig. 2 left panels) and minimum species richness at the site with the least species richness (lower line). As a species occurs at more sites (moving to the right in the figures), the maximum and minimum possible values of average species richness converge on the average of the dataset as a whole. For the BBA datasets, the group of relatively rare species associated with sites of low species richness was dominated by a group of boreal forest species including Canada Jay (*Perisorous canadensis*) and Boreal Chickadee (*Poecile cinctus*). A group of lake and coastal species, such as Common Tern (*Sterna hirundo*), was also associated with sites of low species richness. A number of rare species were associated with sites of high species richness, however. In both BBA datasets, these included a group of wetland species such as American Wigeon (*Anas americana*) and Least Bittern (*Ixobrychus exilis*).

**Figure 4.**
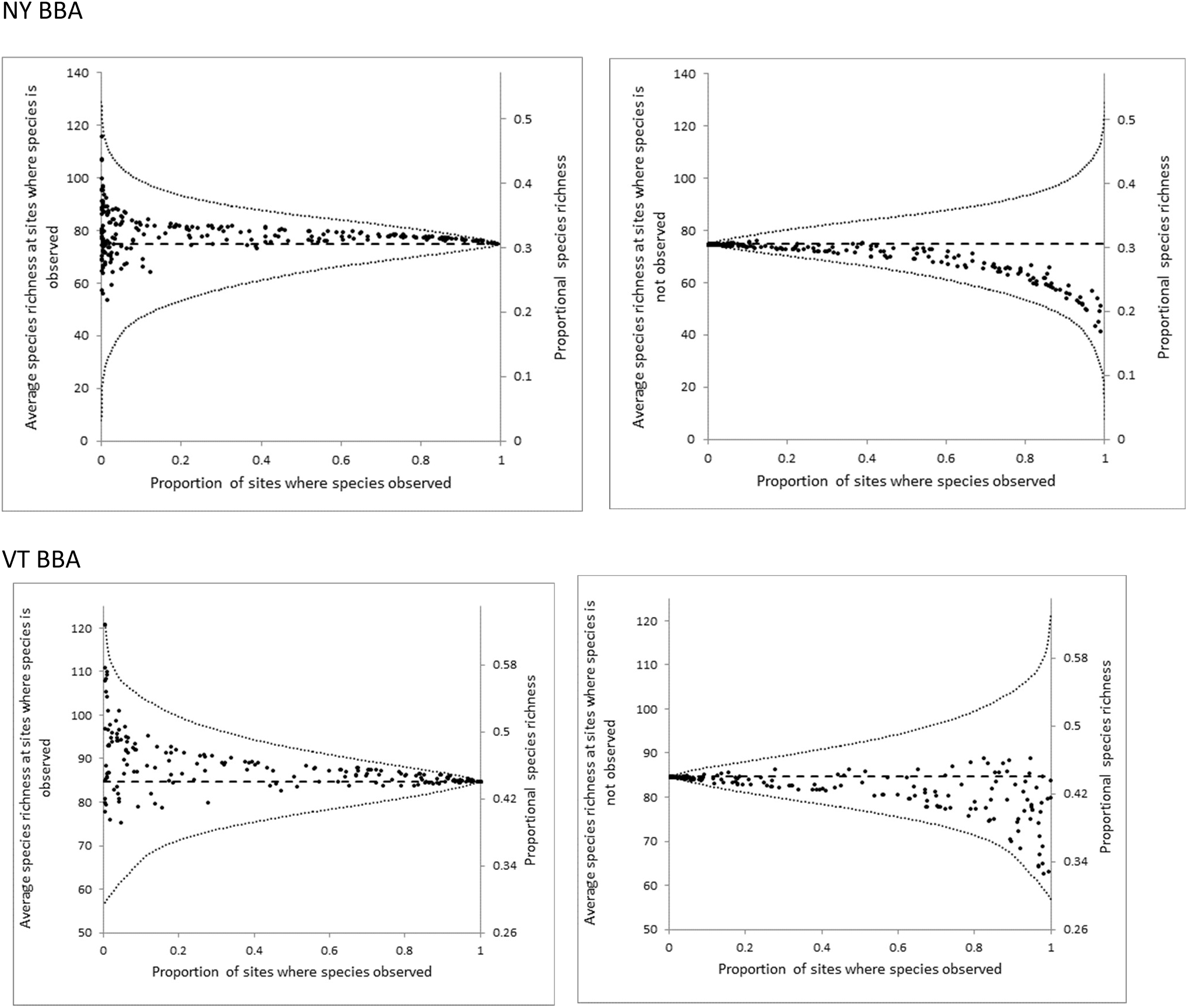
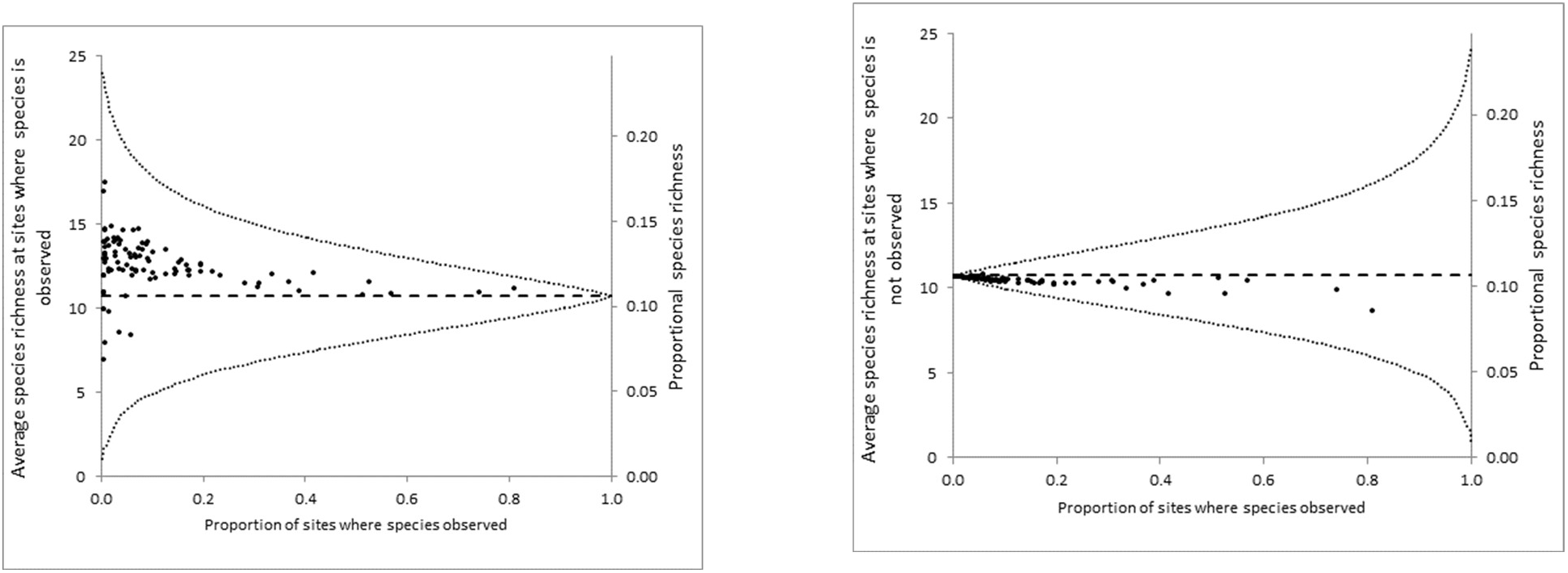
Average species richness at sites where species were recorded (left panels) and where species were <u>not</u> recorded (right panels), as a function of species rarity. Each point represents a single species. Curved dashed lines represent the boundaries within which all points are constrained to occur given the distribution of species richness observed among the sites. Horizontal dashed lines represent average species richness across all sites.

The Vermont point count dataset shared some features with the BBA datasets but also showed differences likely related to spatial extent. Species associated with sites of low species richness included several boreal forest species as in the BBA datasets. They also included several grassland species, such as Eastern Meadowlark (*Sturnella magna*), that were associated with average to above average species richness in the BBA datasets. Grasslands themselves may not have especially high bird species richness, but at the BBA block scale may represent the occurrence of a diversity of land cover types that contribute to species richness. Species in the Vermont point count dataset associated with high species richness tended to use wetland or shrubland habitats; these species were not consistently associated with high (or low) species richness in the BBA datasets.

Unlike the case of examining species richness at sites where species were recorded, when considering sites where species were not recorded (Fig. 4, right panels) common species provided more information about species richness than uncommon species. Mathematical constraints again explain much of the overall patterns of species richness. Rare species by definition are absent from most sites, and therefore the average species richness at those sites approximates that of the overall dataset. Common species are absent from few sites, allowing the potential for greater deviations from dataset average species richness. For the New York BBA and Vermont point count datasets, common species were consistently absent from sites with below average species richness. Most species followed this pattern for the Vermont BBA as well, but the associations were not as strong and some common species were absent from sites with above average species richness. For the Vermont BBA, high species richness was frequently associated with the absence of inhabitants of evergreen or mixed evergreen-deciduous forests.

For both BBA datasets, common species of nonforested habitats were particularly likely to be absent from sites with low species richness. For the Vermont point count dataset, lower species richness was associated with forest interior species (Schwenk and Donovan 2011), such as Black-throated Green Warbler (*Setophaga virens*), and grassland species.

### Species Rarity and Hotspots

Overall, birds of intermediate frequency of occurrence were more likely to be observed in species richness hotspots, relative to non-hotspots, than either the rarest or the most common species. (Fig. 5). The third quartile of mean frequency (i.e., for species of the >50^th^ through 75^th^ percentile) was the highest, and significantly higher than the first quartile, for all three datasets. The second quartile mean was significantly higher than the fourth quartile mean for the Vermont BBA but lower for the other two datasets. That the first quartile would show little difference in frequency within vs. outside of hotspots is largely inevitable from a mathematical standpoint because species occurrence is so low: less than 1% for all species in the Vermont point count and New York BBA datasets and less than 5% for the Vermont BBA. The lower frequencies for the most common quartile of species in the Vermont point count dataset (14-81%) compared to the New York (60-99%) and Vermont (84-100%) BBAs may be an explanation for why this quartile was more associated with hotspots than for the other datasets.

**Figure 5.**
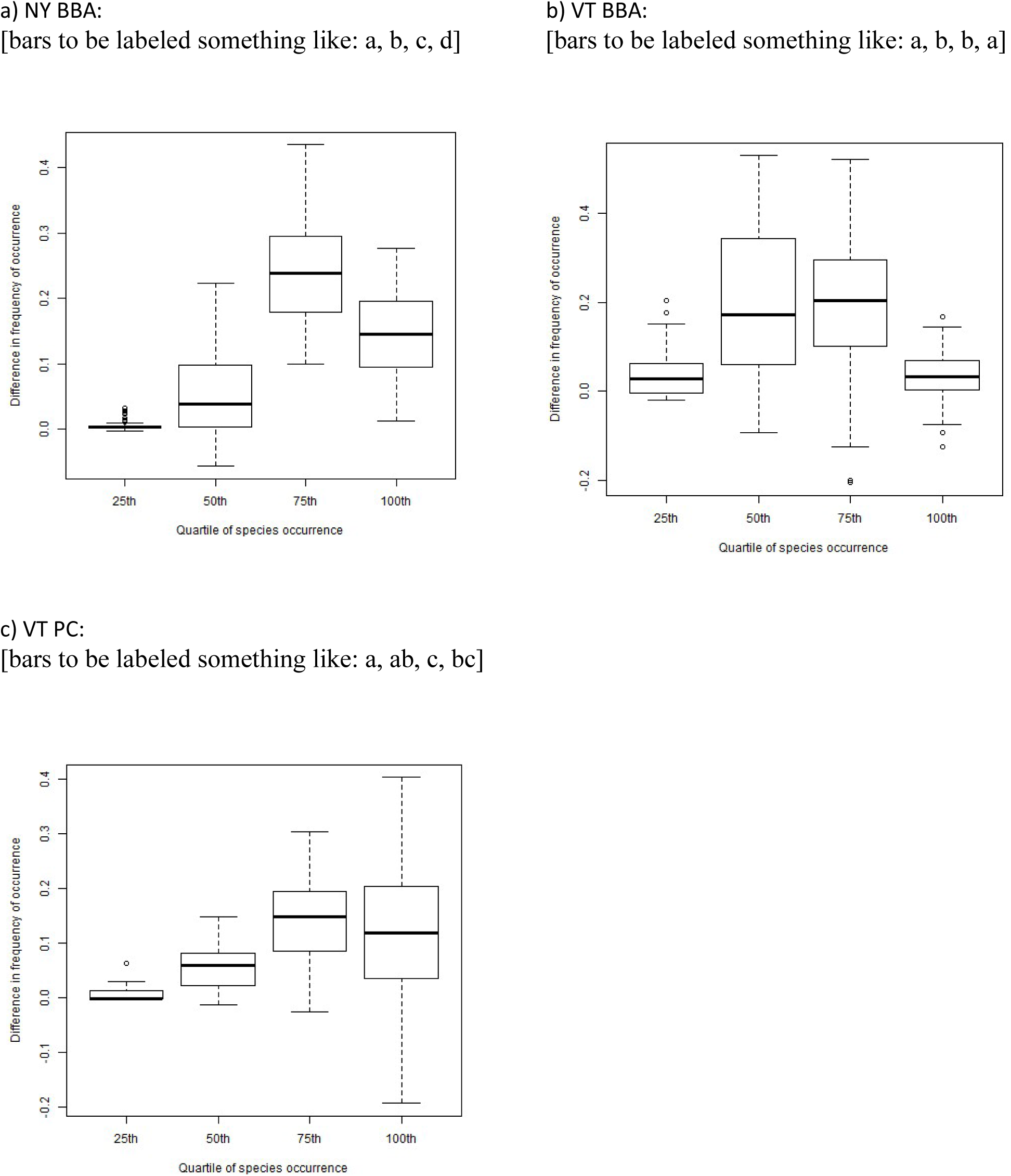
The difference in frequency of occurrence between species richness “hotspots” (10% of sites with highest richness) and the remaining 90% of sites, with each boxplot representing one aggregate of one quartile of species. For example, the quartile labeled “25^th^” refers to the least common species through the 25^th^ percentile of species. Panels: a) New York BBA; b) Vermont BBA; c) Vermont point count dataset.

### Factors Associated with Species Occurrence and Richness

Classifying species into generalized habitat categories was helpful in understanding patterns of species rarity (Table 2). Species primarily depending on ecosystyem types that were uncommon or localized within the study regions, such as wetlands, lakes, and coastal areas, comprised a considerable fraction of species in the less common quartiles of species occurrence. Conversely, species of the abundant deciduous and mixed forest types predominated in the more common quartiles in all three datasets.

**Table 2.** Numbers of bird species recorded in the three surveys by generalized ecosystem type and by quartile.

| Generalized Habitat Type | New York BBA |  |  |  | Vermont BBA |  |  |  | Vermont Point Counts |  |  |  |
| --- | --- | --- | --- | --- | --- | --- | --- | --- | --- | --- | --- | --- |
|  | 25<sup>th</sup><br>"Rare" | 50<sup>th</sup> | 75<sup>th</sup> | 100<sup>th</sup><br>"Common" | 25<sup>th</sup><br>"Rare" | 50<sup>th</sup> | 75<sup>th</sup> | 100<sup>th</sup><br>"Common" | 25<sup>th</sup><br>"Rare" | 50<sup>th</sup> | 75<sup>th</sup> | 100<sup>th</sup><br>"Common" |
| Evergreen forests | 5 | 14 | 9 | 1 | 11 | 6 | 8 | 3 | 4 | 5 | 3 | 3 |
| Deciduous and mixed forests | 4 | 9 | 19 | 26 | 2 | 12 | 8 | 27 | 3 | 6 | 7 | 20 |
| Lakes and coastal areas* | 17 | 8 | 2 | 1 | 8 | 3 | 2 | 0 |  |  |  |  |
| Nonforested uplands | 9 | 8 | 9 | 19 | 7 | 9 | 12 | 8 | 7 | 8 | 7 | 1 |
| Shrublands and young forests | 3 | 5 | 12 | 10 | 5 | 6 | 10 | 7 | 7 | 3 | 7 | 2 |
| Wetlands | 23 | 18 | 10 | 4 | 15 | 12 | 8 | 3 | 4 | 3 | 1 | 0 |

\* Vermont point counts were not designed to survey species using these areas.

Modeling of species occurrence and species richness (Tables 3 and 4) supported these findings and provided additional insight into factors explaining patterns of distributions and diversity. Percent wetlands in a block received substantial support in the BBA datasets but was approximately evenly divided between positive and negative associations with occurrence (Table 3). Positive associations often occurred for relatively uncommon, wetland-dependent species such as Virginia Rail (*Rallus limicola*), whereas negative associations were more often seen for relatively common upland species such as Bobolink (*Dolichonyx oryzivorus*). These associations may have cancelled each other out when it came to species richness, as wetlands had a modest positive effect on species richness in the NY BBA and no significant association for the VT BBA. An analogue for percent wetlands in the Vermont point count dataset, the Topographic Wetness Index, had a similarly mixed effect on species occurrence and a moderately positive association with species richness (Table 4). Elevation was another factor with a mixture of positive and negative associations, particularly in the New York BBA; the association with species richness was negative in Vermont and mixed (depending on ecoregion) in New York.

**Table 3.**
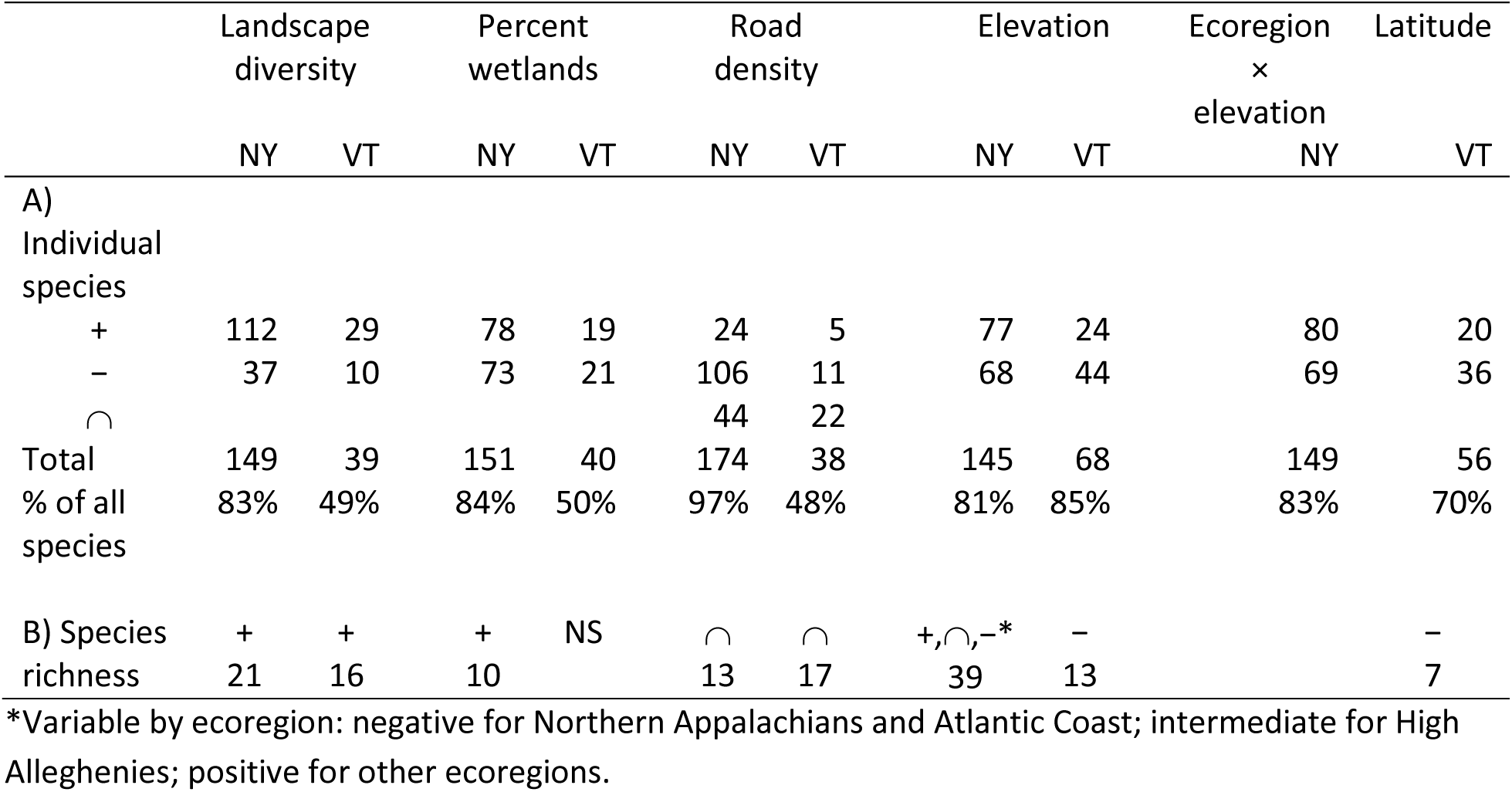
Factors associated with A) species occurrence and B) species richness for the New York (NY) and Vermont (VT) Breeding Bird Atlases. Associations: “+” = positive, “∩” = intermediate or hump-shaped (for factors with a quadratic term), “−” = negative. A) Numbers represent the number of species for which the factor was supported by multimodel selection. For the ecoregion factor, a positive association indicates a species was more likely to occur in the low elevation ecoregions (coastal and Great Lakes). Logistic regression models were developed for 180 NY species and 80 VT species. B) Numbers are a measure of the strength of the effect of the covariate, representing the expected number of species gained from the value of the covariate associated with the minimum species richness to the value associated with the maximum, holding all other covariates constant. NS = inclusion of covariate not supported.

**Table 4.** Factors associated with A) species occurrence and B) species richness for the Vermont point count dataset. Associations: “+” = positive, “∩” = intermediate or hump-shaped, “−” = negative. A) Numbers represent the number of species for which the environmental covariate was supported by multimodel selection. Single-season occupancy models were developed for 67 species. B) Numbers are a measure of the strength of the effect of the covariate, representing the expected number of species gained from the value of the covariate associated with the minimum species richness to the value associated with the maximum, holding all other covariates constant. Inclusion of elevation was not supported.

|  | Road density<br>(within 1 km) | Percent forest<br>(within 1 km) | Percent evergreen<br>(within 300 m) | Distance to edge | Topographic wetness index | Local site forested | Latitude |
| --- | --- | --- | --- | --- | --- | --- | --- |
| A) Individual species |  |  |  |  |  |  |  |
| + | 10 | 21 | 14 | 9 | 6 | 28 |  |
| - | 13 | 9 | 11 | 30 | 8 | 38 |  |
| ∩ | 12 | 24 | 12 |  | 9 |  |  |
| Total | 35 | 54 | 37 | 39 | 23 | 66 |  |
| % of all species | 52% | 81% | 55% | 58% | 34% | —* |  |
| B) Species richness |  |  |  |  |  |  |  |
|  | + | ∩ | + | - | + | + | + |
|  | 2.9 | 3.3 | 1.2 | 7.7 | 1.8 | 1.4 | 2.0 |

\*Included in all models except for one species; *a posteriori* analysis indicated that model selection would have supported selection of the variable for >80% of species.

Other factors were more consistent in their associations with species occurrence and richness. Landscape diversity had a predominantly positive association with species occurrence and a strong positive association with species richness in both BBA datasets (Table 3). The association cut across all degrees of rarity but was especially prevalent among common species. An analogue to landscape diversity at the point count scale, distance to edge of another ecosystem type, showed similar patterns. Many more species had a negative association with distance to edge (i.e., they were more likely to be observed closer to edges) than a positive one and this factor was strongly negative in the species richness model for the Vermont point counts (Table 4).

The influence of human development on species occurrence and richness was evident through the road density covariate (Tables 3 and 4). In New York, which has a much greater density of roads than Vermont, more than four times as many species had a negative association with roads than a positive one. In heavily forested Vermont, an intermediate association was most frequent in the BBA and associations were mixed among species in the point count dataset. Forest species previously identified as being area-sensitive (e.g., Robbins, Dawson, and Dowell 1989), such as Scarlet Tanager (*Piranga olivacea*), frequently showed negative associations with road density. Many of the positive or intermediate associations between occurrence and road density occurred among open and edge species such as American Crow (*Corvus brachyrhynchos*) and Mourning Dove (*Zenaida macroura*). Reflecting this diversity of responses to road density, species richness peaked at intermediate levels of road density in both BBA datasets; at the Vermont point-count scale, the association with species richness was moderately positive.

## Discussion

While I found that in some situations common species convey more information about species richness patterns than rare species, from conservation and practical standpoints they have limitations. The cautions I present against solely relying upon common species as proxies for species richness may to serve temper more sweeping assertions of their applicability (Lennon et al. 2004, Mazaris et al. 2008). As appealing as shortcuts to assessment are, there are applications where they cannot substitute for thorough surveys of the full spectrum of rare to common species.

I did find that based on sequential correlation of partial species sets, correlations approached unity with total species richness more rapidly beginning with common species than with rare species at both spatial extents and resolutions. Šizling et al. (2009) have demonstrated based solely on mathematical considerations that this pattern is to be expected when the number of rare species substantially exceeds the number of common species. Such frequency distributions are common in species richness studies (Ulrich and Ollik 2004). I also found, however, that random sequences of species sometimes outperformed the ordered sequence beginning with the most common species. Therefore, just because common species explain more of the overall pattern of species richness than rare species at a particular spatial scale, one should not assume that a set of the most common species *best* explains patterns of species richness.

I also found that species occuring with intermediate frequency, rather than the most common (or most rare) species, tended to be most associated with “hotspots” of species richness. Understanding the locations of hotspots may be particularly important from a conservation standpoint (Reid 1998). Where I found common species to be most informative was in the identification of “coldspots;” the absence of common species often signaled that overall species richness was low. Such areas might be lower priorities for conservation action. However, they should be examined more closely to ensure that low observed species richness is not simply a function of depressed survey effort or detection rates. Moreover, I found that some rare species of conservation concern in the study areas are associated with areas of low species richness.

The differences I found across spatial scales are consistent with previous work (Hess et al. 2006, Rahbek 2005, McKerrow et al. 2018) and serve as a further caution against uncritically accepting the premise that common species determine the majority of patterning in species richness. Although species relatively common at a fine-grain size (point count circles, 1.8 ha) tended to also be common at a coarse-grain size (atlas blocks, 25 km^2^), the actual frequencies varied dramatically and the sequence of which species were most frequent also varied. Also, many of the rarest species at the coarse-grain size were not detected at all at the fine scale. Furthermore, the degree to which the occurrence patterns of a species was correlated with species richness at one spatial resolution was minimally predictive of its correlation at another spatial resolution. As a consequence, the information and conclusions that can be drawn from a set of species common (or rare) at one scale in this study are not consistent with what that set conveys at a different spatial scale.

Understanding the factors that influence species distributions, and how they vary among common and rare species, can help in explaining the ability of common species to capture patterns of species richness. Rare species have been reported to differ from common species based on factors such as having lower dispersal ability, larger body size, higher trophic level, use of less common resources, and differential associations with environmental variables (Lennon et al. 2011, Gaston and Kunin 1997). In this study, rare species appear to largely be those that specialize on relatively uncommon ecosystem types such as seacoasts and boreal forests or otherwise preferentially occur under climate regimes that are peripheral to the study areas. Many of the most common species inhabitated the study areas’ matrix deciduous forests and were at least moderately tolerant of human activities. Comparing the two spatial resolutions of the study, species richness tended to accumulate through inclusion of adjacent areas supporting different sets of species rather than by encountering areas particularly suited to simultaneous, overlapping use by species. Therefore, heterogenous landscapes – associated with increased diversity in this and other studies (e.g., Böhning-Gaese 1997, Lepczyk et al. 2008) – probably provide sufficient matrix habitat for the common species while offering varied other habitats that allow multiple rarer species to occur as well. From this perspective, it is apparent why the presence of common species would not be a good indicator of high species richness. The absence of common species by contrast could suggest landscapes dominated by land cover uncharacteristic of the region as a whole, such homogenous urban or agricultural areas with relatively low species richness.

In summary, I found little evidence that common species alone could provide a satisfactory shortcut to understanding biodiversity, particularly given that rare species are often the ones facing the greatest risk of extirpation or extinction. Common species may capture a substantial amount of spatial patterning, but rare and intermediate-frequency species each offer unique information and advantages as well. Proxies may be most helpful in situations with very large numbers of species that are relatively difficult to survey in comparison to potential proxies, which generally was not the case in the avian examples considered here. Drawing upon citizen scientists to aid in monitoring and ensuring that unusual or unique ecosystem types and configurations are surveyed may be invaluable in obtaining the thorough understanding of biodiversity needed for successful conservation outcomes.

## Acknowledgments

I greatly appreciate assistance, reviews, and data provided by Therese Donovan (Vermont point count dataset) and Rosalyn Renfrew (Vermont Breeding Bird Atlas). The Vermont Breeding Bird Atlas was made possible by the participation of hundreds of volunteer participants and numerous sponsors including the Vermont Center for Ecostudies and the Vermont Fish and Wildlife Department. The New York Breeding Bird Atlas was also supported by hundreds of volunteer participants as well as the New York State Department of Environmental Conservation and many other sponsors. The Vermont point count dataset was made possible by surveys conducted by T. Collingwood, R. DeMots, C. Eiseman, and S. Wilson, access granted by many landowners, and funding provided by the U.S. Department of Agriculture Northeastern States Research Cooperative, the McIntire-Stennis Cooperative Forestry Program, the U.S. Forest Service, and the U.S. Geological Survey, Vermont Cooperative Fish and Wildlife Research Unit.

